# Histochemical Examination of Changes In Testis After Vasectomy In Rats And Qualitative Analysis Of Tissue Zinc Content

**DOI:** 10.1101/2022.01.21.477201

**Authors:** Emine Ozcinar, Yasemin Ersoy Canillioglu, Sule Cetinel

**Affiliations:** Department of Histology and Embryology, School of Izmir Tinaztepe University, Izmir, Turkey; Department of Histology and Embryology, School of Medicine, Bahcesehir University, Istanbul, Turkey; Department of Histology and Embryology, School of Medicine, Marmara University, Istanbul, Turkey

**Keywords:** vasectomy, zinc, testis, TEM, EDS

## Abstract

It is claimed that around 100 million men worldwide have undergone vasectomy for birth control. Vasectomy is an operation to prevent the transfer of sperm by preventing the luminal continuity of the vas deferens. Degenerative changes after vasectomy usually start with loss of spermatogenic cells in the germinal epithelium, while Sertoli cells are affected later. There is no change in Leydig cells. Zinc, which is found at high rates in the male reproductive system, is an element in the structure of many enzymes necessary for cell membrane integrity, growth and sexual maturity.

In this study, bilateral vasectomy was performed on 18 young adult male rats and control operation was performed on 18 rats. Afterwards, 6 animals from each group were sacrificed after 6 weeks, 12 weeks and 12 months. Germinal epithelium, tubule structure, basement membrane, interstitium and collagen fiber changes were examined at microscope and transmission electron microscope level. On the other hand, zinc content in testicular tissue was measured by Energy-dispersive X-ray spectroscopy (EDS). Six weeks after vasectomy, localized losses in the germinal epithelium and disorganization of spermatogenic cells were observed. In addition to the loss of spermatogenic cells at the 12th week, separations, thickening and an increase in the collagen fiber in the peritubular area were observed in the interstitium and perinuclear area. Tissue zinc content was also found to be the lowest in the 12 months group.

In conclusion, our study showed that the degenerative changes in the rat testis that increased with time after vasectomy were parallel to the decrease in tissue zinc content.

## 1. INTRODUCTION

Vasectomy is a family planning method that is used safely in male reproductive control and is increasingly preferred today. Therefore, researchers have a keen interest in examining the results of vasectomy. With the vasectomy operation, it is aimed to achieve maximum safety sterility and minimal post-operative complications. Many techniques are used in vasectomy operations, regardless of the technique used, 97.2-99% success is achieved in birth control. The most common short-term complications of vasectomy are hematoma, infection, and sperm granuloma formation. In the long term, vasitis nodosa, chronic pain and even prostate cancer can be counted. In animal studies conducted to investigate the sequelae of vasectomy, testicular degeneration was found to increase in direct proportion to the time, while significant testicular degeneration findings were not detected in human studies. The reasons for this have not been clarified yet (1–4). After vasectomy, the testis continues to produce sperm. Since the vas deferens is attached, the area they are in is limited for these produced sperms. Therefore, the intraluminal pressure rises, especially in the epididymis. Due to anatomical differences, this pressure is transmitted to the seminiferous tubules in a short time in some animals. As the time increases, the epididymis loses its integrity from place to place and sperm granulomas form. In humans, however, due to the easy tearing of the epididymis due to the increase in pressure in the epididymis, high pressure does not reflect on the seminiferous tubules, but sperm granulomas form in a short time, which is thought to balance the pressure (1,5–7).

Zinc is an essential element for all living things and is present in more than 100 enzymes. High levels of zinc in the male reproductive system have been shown to significantly affect sperm motility, but its mechanism has not been fully clarified (8,9). It is thought that zinc has a direct stimulating effect on steroid synthesis and especially on testosterone synthesis (10). In recent years, the detection of the mRNA of ZnT-3, the zinc carrier protein, in the testis has added a new dimension to the studies on zinc in male fertility (11).

In many studies conducted to date, testicular tissue after vasectomy has been examined, but elemental zinc analysis of testicular tissue has not been performed. Our aim with this study is to examine the early and late morphological changes in the testis after vasectomy and to question the relationship between tissue zinc content and these changes.

## 2. MATERIAL AND METHODS

The study was initiated with the approval of Marmara University Experimental Animals Local Ethics Committee. In this study, a total of 36 male Wistar albino rats (230-410 grams), obtained from the laboratory of experimental animals of Marmara University, were used. Animals were fed standard rat chow and drank tap water in appropriate cages for 12 hours in light-dark conditions.

2 experimental groups were formed; bilateral vasectomy was performed on 18 rats in the vasectomy group under general anesthesia (Rompun 2%, Ketalar 50mg/ml; 210μl Ketalar and 90μl Rompun intraperitoneal). The abdomen was opened by passing the skin and abdominal wall with a 2 cm long transverse incision 2 cm above the penis. The testis, vas deferens and epididymis were taken out. By placing forceps under the vas deferens, knots were tied with a silk thread at an interval of approximately 1 cm on both ends of the canal between the tips of the forceps, and the part in between was cut and removed. The same procedure was done on the other side. Afterwards, the organs were placed in the abdomen and the abdomen was sutured. At the end of 6 weeks, 12 weeks and 12 months, 6 animals were sacrificed. Eighteen rats in the control group were operated under general anesthesia by opening and closing the abdomen, and the same number of animals were sacrificed at the same time as the vasectomy group, and testicular tissues were removed.

Testicular tissue samples from all groups were fixed with 10% neutral formalin for 18 hours. Tissues were dehydrated by passing through increasing alcohol series and then cleared with toluene. After being kept in paraffin for 1 night in an oven at 60°C, blocks were formed by embedding in fresh paraffin at room temperature. Afterwards, 5μm thick sections were taken from these blocks and hematoxylin-eosin staining was performed for general histological evaluation, and Von-Gieson staining was performed to evaluate collagen fiber amount and changes. Stained sections were examined with an Olympus BH2 microscope. For fine structure examination, testicular tissue samples from all groups were fixed in 2.5% 0.1 M phosphate buffered (pH 7.2) glutaraldehyde fixative at 4°C for 4 hours. Post-fixation was performed for 1 hour with 1% osmium tetroxide prepared in the same phosphate buffer. Afterwards, the tissues were passed through increasing alcohol series and propylene oxide and embedded in epon 812. Semi-thin sections taken in an ultramicrotome (Leica Ultracut R) were stained with toluidine blue. In these sections, thin sections with a thickness of approximately 60 nm taken from the areas containing seminiferous tubules were placed on copper grids and contrasted with uranyl acetate and lead citrate (12). Contrasted grids were examined with JEOL 1200 SX TEM.

For tissue zinc analysis, testicular tissue samples from all groups were fixed in 2.5% 0.1M Phosphate buffered (pH 7.2) glutaraldehyde at 4°C for 4 hours. Post-fixation was performed for 1 hour with 1% osmium tetroxide prepared in the same phosphate buffer. Afterwards, the tissues were passed through increasing alcohol series and passed through 2/1, 1/1, ½ alcohol and amyl acetate series, respectively, and were taken into pure amyl acetate. In the elemental analysis of tissues with EDS, the qualitative ratios of the main elements of the tissue, carbon (C), oxygen (O), osmium (Os) used during preparation, gold (Au) used in tissue coating, and zinc (Zn), the amount of which we investigated, were compared.

After 6 weeks, 12 weeks and 1 months, 1 section was taken transversely from the testis tissue of all animals sacrificed and 5 μm thick sections were taken, skipping 5 sections. From the sections stained with hematoxylin-eosin, the properties of a total of 100 tubules for each animal were evaluated, with 10 tubules in each section, at X200 magnification under the light microscope, and their arithmetic averages were taken. In the examined seminiferous tubules, smooth tubule contours, regular placement of spermatogenetic cell lines and appropriate height, and normal thickness of the basement membrane and interstitial area were taken as criteria for tubule with normal morphology (3,9).

## 3. RESULTS

The results of the semi-quantitative analysis of the macroscopic appearances of testicular tissues and seminiferous tubules of all groups are shown in Table 1.

**Table 1.**
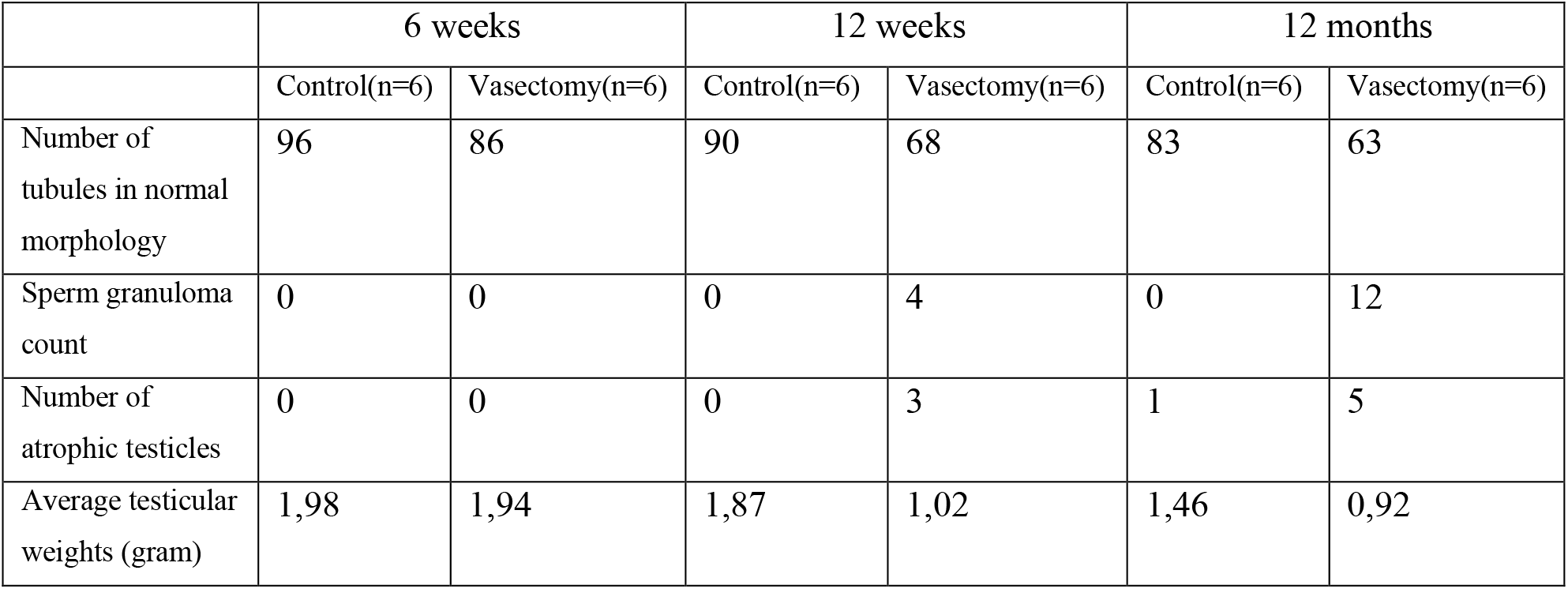
Semiquantitative analysis of testis macroscopic findings and seminiferous tubules

|  | 6 weeks |  | 12 weeks |  | 12 months |  |
| --- | --- | --- | --- | --- | --- | --- |
|  | Control(n=6) | Vasectomy(n=6) | Control(n=6) | Vasectomy(n=6) | Control(n=6) | Vasectomy(n=6) |
| Number of tubules in normal morphology | 96 | 86 | 90 | 68 | 83 | 63 |
| Sperm granuloma count | 0 | 0 | 0 | 4 | 0 | 12 |
| Number of atrophic testicles | 0 | 0 | 0 | 3 | 1 | 5 |
| Average testicular weights (gram) | 1,98 | 1,94 | 1,87 | 1,02 | 1,46 | 0,92 |

Control groups, hematoxylin and eosin stainings at 6 (Figure 1A) and 12 (Figure 1B) weeks, testes were found to have normal morphological structure. Spermatogenic cells were arranged regularly in the seminiferous tubules. Many mature sperm were observed in the lumen. The tubule contours were smooth and the interstitial area and Leydig cells were normal. In the testicular tissues obtained at the end of 12 months (Figure 1C), a decrease was found in the number of mature sperm in the lumen compared to the other control groups. Ondulation in tubule contours and thickening in interstitium were observed.

**Figure 1.**
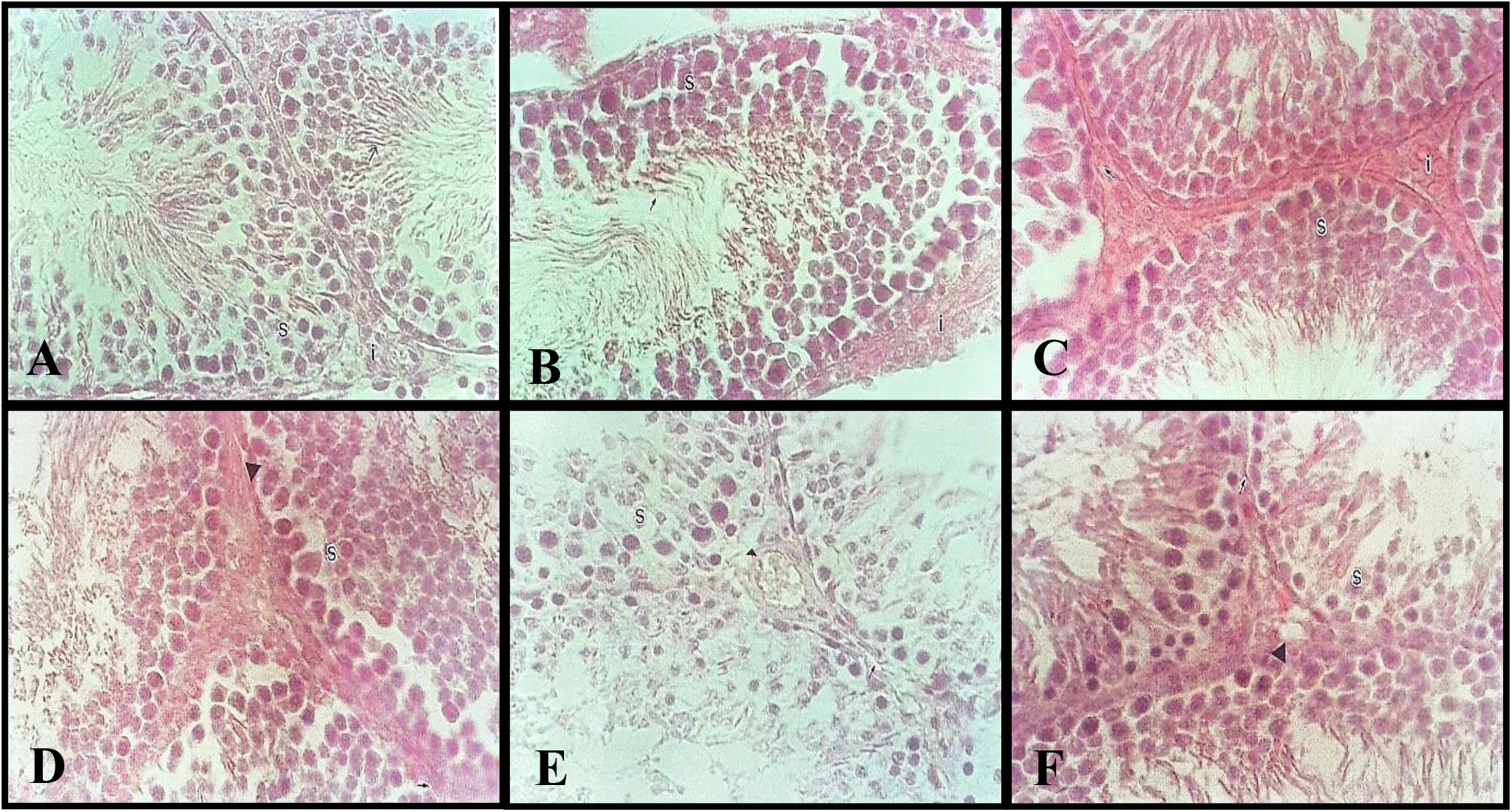
Testicular hematoxylin-eosin staining

Von Gieson staining performed to show collagen fibers showed that the amount of collagen in the basement membrane and interstitium in testis tissues increased at 12 months (Figure 2C) compared to 6 weeks (Figure 2A) and 12 weeks (Figure 2B).

**Figure 2.**
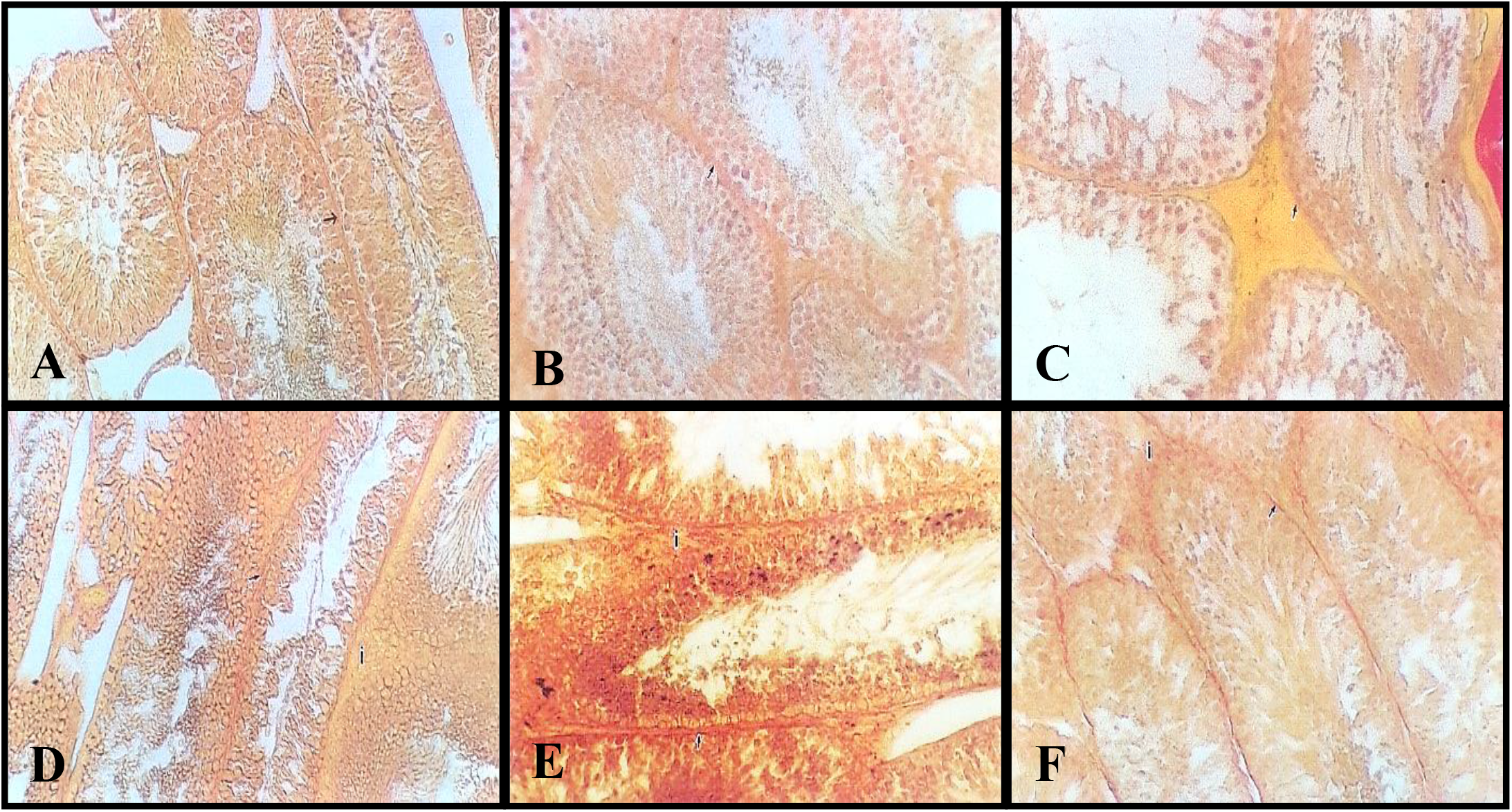
Testis Von-Gieson staining

As a result of staining the semi-thin sections with Toluidine Blue, which has metachromatic properties, the cells and basement membrane structure in the testicular tissues of the control groups were normal at 6 (Figure 3A), 12 weeks (Figure 3B) and 12 months (Figure 3C).

**Figure 3.**
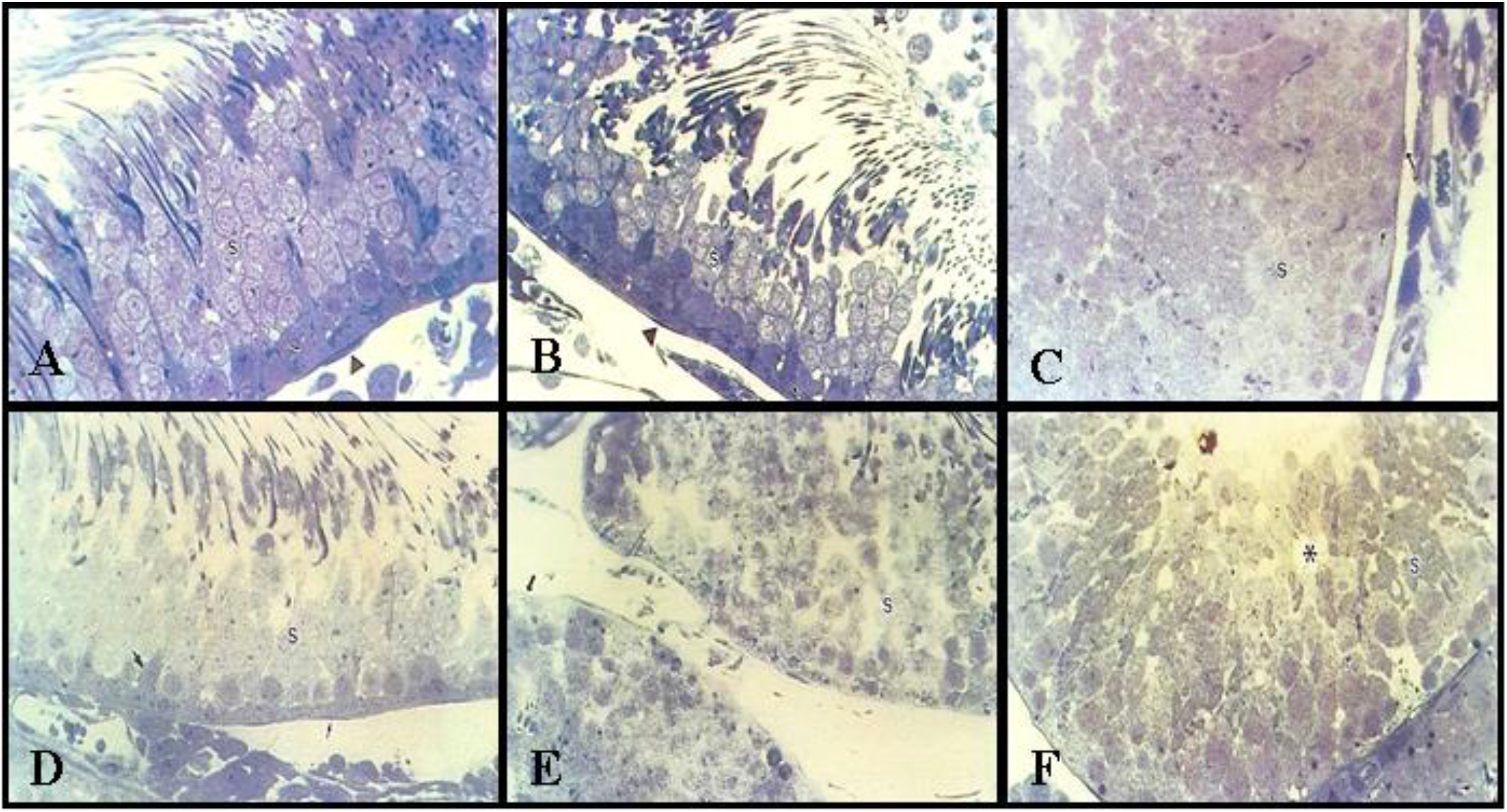
Testicular toluidine blue staining

In the examinations performed with TEM, the basement membrane was in normal structure at 6 and 12 weeks. Sertoli cells and fine structure of myoid cells were in normal morphology. The organization of spermatogenic cells was normal and there was a normal amount of collagen fibers in the tubule walls. Animals at the end of 12 months showed an enlargement of the area between Sertoli and myoid cells and a slight increase in the amount of collagen.

The comparative analysis of zinc and other elements in all groups with EDS is shown in Table2.

**Table 2.** Tissue zinc and element determination with EDS

|  | 6 weeks |  | 12 weeks |  | 12 months |  |
| --- | --- | --- | --- | --- | --- | --- |
|  | Control(n=6)% | Vasectomy(n=6)% | Control(n=6)% | Vasectomy(n=6)% | Control(n=6)% | Vasectomy(n=6)% |
| C(Carbon) | 29.05 | 42.51 | 40.69 | 35.46 | 47.28 | 36.12 |
| O(Oxygen) | 15.24 | 24.08 | 25.53 | 23.63 | 29.44 | 24.26 |
| S(Sulfur) | 0.21 | 1.28 | 0.68 | 0.43 | 0.49 | 0.38 |
| Zn(Zinc) | 0.86 | 0.16 | 1.32 | 0.39 | 1.88 | 0.13 |
| Os(Osmium) | 10.14 | 5.75 | 6.86 | 14.04 | 4.95 | 16.00 |
| Au(Aurum) | 44.51 | 26.22 | 24.91 | 26.04 | 15.96 | 23.11 |

Vasectomy groups, Hematoxylin-eosin staining at 6 weeks (Figure 1D) showed minimal irregularities in the arrangement of spermatogenic cells, occasional undulations in tubule contours, and minimal separations in interstitial areas. The number of tubules without mature sperm was higher than in the control group. At 12 weeks (Figure 1E), it was determined that the irregularity in the arrangement of spermatogenic cells increased and their number decreased, the corrugation in the tubule contours became more pronounced and the separations in the interstitium became more pronounced. At 12 months (Figure 1F), the number of tubules without mature sperm was found to be increased compared to the 6- and 12-Week vasectomy groups. In addition, the separations in the interstitium became quite evident.

In Van Gieson staining, the amount of collagen in the basement membrane and interstitium was found to be higher in the group at the end of 12 months (Figure 2F) compared to the 6- (Figure 2D) and 12-week (Figure 2E) vasectomy groups. In the examinations performed with toluidine blue, a decrease in the number of germinal epithelium and spermatogenic cells and deteriorations in organization and sequence were detected at 6 weeks (Figure 3D). Sertoli and myoid cells were normal. At 12-weeks (Figure 3E) and 12 months (Figure 3F), additional corrugation was observed in intraepithelial vacuoles and tubule basement membrane.

In the examinations performed by TEM, very slight separations between Sertoli cells and myoid cells and some spermatogenic cells and a small number of vacuoles were detected in the cytoplasm of Sertoli cells at 6-weeks. An increased amount of collagen fiber was detected in the area between Sertoli cells and myoid cells compared to the vasectomy and control groups for 6-weeks. In 12 months, a significant increase in vacuoles in the cytoplasm of Sertoli cells, in addition, a prominent increase in the nucleolus and a significant increase in the amount of basement membrane collagen were detected.

## 4. DISCUSSION

The effect of vasectomy on the human testis is not yet fully understood. In many studies, a decrease in testicular weight up to atrophy was observed depending on the time elapsed after vasectomy (13, 14–16). One study found a significant ipsilateral weight reduction 4 weeks after unilateral vasectomy. In this study, it was thought that the cause of atrophy was fibronectin and type 1 collagen secreted from myoid cells (13). In another study, 4 weeks after bilateral vasectomy, both testicular weights were decreased, 6 months later, weight loss continued and significant testicular atrophy developed in some of the rats, and an increase in the number of apoptotic cells was shown. It has been suggested that the reason for all these degenerative changes is increased intraluminal pressure and apoptosis developing accordingly (14). In another study, 50% testicular atrophy was detected 6 months after unilateral vasectomy in rats, and it was observed that atrophy was bilateral in some rats. In this study, immunological causes were considered as the cause of atrophy (16). In parallel with the studies performed in our study, a minimal decrease was detected in testicular weight compared to the control group 6 weeks after vasectomy, and it was observed that the decrease continued as the time progressed. It was thought that this situation may be due to the increase in hydrostatic pressure in the lumen since it is dependent on the vas deferens.

In the studies performed, sperm granulomas were found in varying numbers and localizations in the epididymal canal on the vasectomy side following vasectomy in rats. These are observed as early as 6 weeks and their number increases over time. The increase in pressure in the epididymis and the resulting chronic inflammatory reaction have been shown to be the cause of granulomas (5,13,14,16). In our study, 12 weeks after vasectomy, 1 out of 6 rats had bilateral sperm granulomas and 2 had unilateral sperm granulomas. Bilateral sperm granulomas were observed in all rats 12 months after vasectomy. There was no one-to-one relationship between the development of granuloma and testicular atrophy.

In a study, a decrease in germinal cells was detected in both testicular seminiferous tubule epithelium 4 weeks after unilateral vasectomy, while Sertoli cells were observed to be normal. In this study, serum reactive oxygen radicals were found to be high. This has been associated with testicular damage after vasectomy (4). In another study, localized loss of germinal epithelium and decreased cell number in tubules were found 12 weeks after bilateral vasectomy. After 6 months, in addition to these findings, enlargement in the interstitial area and an increase in the amount of collagen were observed (3). In our study, it was observed that the irregularity in the spermatogenic cell arrangement in the seminiferous tubules, which started 6 weeks after vasectomy, continued after 12 weeks in the form of significant organizational disorder and a decrease in cell numbers. Ondulation in tubule contours and separations in interstitium were noted. No change was detected in Leydig cells. In the examinations performed 12 months after vasectomy, it was observed that all these degenerative changes increased.

In one of the studies, an increase in the amount of collagen in the peritubular area and irregularities in the Sertoli cell nucleus were noted in the examination performed with TEM 4 weeks after vasectomy (10). In another study, extracellular spaces were found in the germinal epithelium after 12 weeks, and basement membrane thickening after 6 months, and nuclear openings and chromatin condensation in spermatogenic cells were found (3). In our study, a small number of vacuoles and minimal separations between spermatogenic cells were observed in Sertoli cell cytoplasm 6 weeks after vasectomy in the TEM examination, and these findings increased 12 months later. In addition, there was an increase in the amount of collagen in the area between Sertoli and myoid cells. These findings were thought to be the result of degeneration that increased over time.

In some studies, in humans, it has been determined that zinc ion affects sperm motility, but its mechanism has not been fully explained. In one of the studies, serum zinc levels were found to be low in the infertile group, but no difference was found between the fertile and infertile groups in terms of semen zinc levels (8). In another study, serum and semen zinc levels were found to be lower in infertile people compared to the fertile group (17). In another study, it was shown that zinc is found extensively in spermatogonia, spermatocytes and spermatids in rat testicles (11). In our study, qualitative zinc measurement was performed in the tissue to compare the changes in the testis after vasectomy with the tissue zinc ion. When each group was compared with its own control group, it was observed that the zinc content of the testis tissue began to decrease 6 weeks after vasectomy, and the difference between the control group and the control group increased over time.

## 5. CONCLUSION

It was observed that degenerative changes developed in the testis, especially in spermatogenic cells, in direct proportion to the time passed after vasectomy in rats. Parallel to these degenerative changes, there was also a significant decrease in the zinc content of the testis tissue. In the light of previous studies and the findings we obtained, it was thought that low zinc levels may play a role in the increase of degenerative changes in the testis and that the decrease in tissue zinc content detected by the EDS method after vasectomy may be the result of cell loss as a result of degeneration.

## 7. ACKNOWLEDGEMENTS

The authors would like to thank Associate Professor Fatih Oltulu and Assistant Professor Gurur Garip for data analysis.

